# SARS-CoV-2 B.1.617 Indian variants: are electrostatic potential changes responsible for a higher transmission rate?

**DOI:** 10.1101/2021.06.08.445535

**Authors:** Stefano Pascarella, Massimo Ciccozzi, Davide Zella, Martina Bianchi, Francesca Benetti, Francesco Broccolo, Roberto Cauda, Arnaldo Caruso, Silvia Angeletti, Marta Giovanetti, Antonio Cassone

**Author notes:** Denote equal contribution (Stefano Pascarella, Massimo Ciccozzi).

## Abstract

Lineage B.1.617+, also known as G/452R.V3, is a recently described SARS-CoV-2 variant under investigation (VUI) firstly identified in October 2020 in India. As of May 2021, three sublineages labelled as B.1.617.1, B.1.617.2 and B.1.617.3 have been already identified, and their potential impact on the current pandemic is being studied. This variant has 13 amino acid changes, three in its spike protein, which are currently of particular concern: E484Q, L452R and P681R. Here we report a major effect of the mutations characterizing this lineage, represented by a marked alteration of the surface electrostatic potential (EP) of the Receptor Binding Domain (RBD) of the spike protein. Enhanced RBD-EP is particularly noticeable in the B.1.617.2 sublineage, which shows multiple replacements of neutral or negatively-charged amino acids with positively-charged amino acids. We here hypothesize that this EP change can favor the interaction between the B.1.617+RBD and the negatively-charged ACE2 thus conferring a potential increase in the virus transmission.

## Introduction

The Coronavirus Disease 19 (COVID-19), caused by the new coronavirus SARS-CoV-2, continues to spread worldwide, with more than 163 million infections and about 3,5 million deaths as of May 17^th^, 2021 (www.who.int). In order to fight this dreadful disease, several safe and efficacious vaccines against SARS-CoV-2 are being used with remarkable effectiveness in some countries and limited availability in others. In particular, the capacity of some countries to halt SARS-CoV-2 spread is still limited due to inadequate resources and vaccination infrastructures (1,2). In this scenario, several SARS-CoV-2 variants have been identified and have become a global threat. Some of them have been classified as variants of concern (VOCs) due to their mutations in the S1 subunit of the spike (S) protein, particularly in its receptor binding domain (RBD) (3–5). One of them, identified as B.1.1.7, also known as the UK variant, bears a substitution of asparagine with tyrosine on the position 501 and a deletion of two amino acids in the position 69-70 of the S1 subunit. This variant has quickly spread in several European countries to become globally dominant (5). Others VOCs have been isolated in South Africa and Brazil and have been studied for their enhanced contagiousness and resistance to neutralization by antibodies from convalescent and vaccine-recipient subjects (6–8) (**Table 1**). Quite recently, a new variant under investigation (VUI) has been isolated from Maharashtra, India, in a setting of highly diffusive epidemic with devastating proportions (. This variant, identified as B.1.617 carries several non-synonymous mutations. Two of them, the E484Q (or the P478K) and the L452R, are located in the RBD region, and they are critical site for the binding with ACE2. Initial data suggest these mutations could confer increased transmission and immune-evasion (9–11). Currently, B.1.617 comprises three subvariants, B.1.617.1-3, with different distribution of the mutations P478K and E484Q (Public Health England). Here we focus on biochemical and biophysical changes conferred to the B.1.617+ VUI by the P478K and E484Q mutations. We then compare these changes with other VOC in order to establish whether and to what extent those amino acid changes can influence the interaction of the Spike protein with ACE2, thus potentially affecting SARS-CoV-2 transmission and immune-escape properties.

**Table 1.**
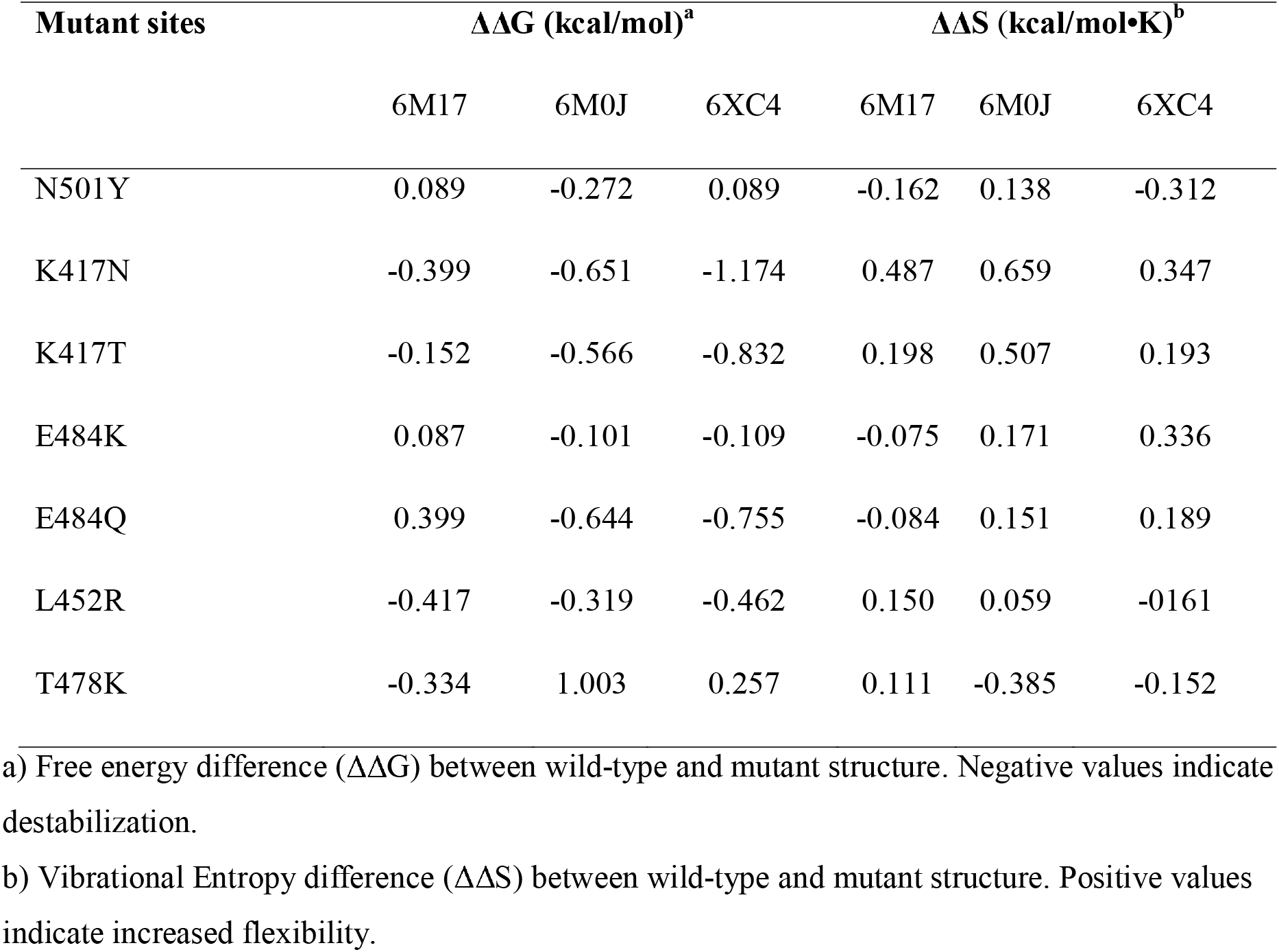
Dynamut results

## Methods

In order to perform more robust analysis 3 different 3-dimensional structures of SARS-CoV-2 Spike Glycoprotein have been downloaded from Protein Data Bank with the following characteristics:

1. SARS-CoV-2 RBD in complex with neutralizing antibody CC12.3 (PDB code: 6XC4 chain A)
2. SARS-CoV-2 Spike Glycoprotein whole structure (PDB code: 6XR8 chain E)
3. SARS-CoV-2 RBD in complex with ACE2-B0AT1 (PDB code: 6M17 chain F).

The DynaMut server (12) has been used to predict the impact of mutations on protein stability analyzing the folding free energy (ΔΔG) and the Vibrational movement (ΔΔS), two crucial characteristics of the function and the molecular recognition of the protein. PyMol (PyMol, version 2.4) suite was utilized for in-silico mutagenesis and its Adaptive Poisson–Boltzmann Solver (APBS) plugin (13) has been used to calculate the electrostatic potential of the wild-type and VOCs SARS-CoV-2 Spike Receptor Binding Domain (RBD) and evaluate potential differences in terms of molecular interaction with ACE2 receptor. The results have been reported within a range between −5 kT/e and +5 kT/e.

## Results

### Protein stability

In silico prediction of the mutation impact on the RBD stability has been carried out with DynaMut. Three alternative RBD structures denoted by the PDB codes 6M17, 6M0J and 6XC4 have been tested. These structures display small differences in the conformation of loops, especially in the one inside the Receptor Binding Motif (RBM). According to the parameters of our in silico experiments, the output of DynaMut for mutant sites within loops differ depending on the starting loop conformation. For this reason, we used a normalized procedure, whereby the same mutation has been tested in each of the three different RBD structures. The effect on stability, de- or stabilization, has been evaluated following a majority criterion. Detailed results have been reported in **Table 1**.

Starting with position 501 within a loop in the RDB interacting with ACE2 receptor, we note that the mutation N501Y does not show any unambiguous structural effect and should be considered neutral (**Table 1**). Also, position 417 is within the interface α-helix where Lys interacts with ACE2 Asp30. However, in this case, our results predict that both mutants (K417N and K417T) could destabilize the protein and increase local flexibility. Similarly, Glu484 is in an interfacial loop and interacts with ACE2 Lys31. Again, our data predict that both mutations (E484K and E484Q) exert a destabilizing effect, with E484Q being the strongest. We note that Glutamic Acid is a polar, negatively charged, hydrophilic amino acid while Lysine is basic, chiral, charged and aliphatic amino acid and its ε-amino group often participates in hydrogen bonding, salt bridges and covalent interactions. Both mutants E484K and E484Q, would likely increase local molecular flexibility. L452 is in a short β-strand and it is exposed to the solvent. Apparently, it does not interact directly with ACE2. Mutation L452R is predicted to be destabilizing with increased local flexibility. Position 478 is in a loop in proximity of the ACE2 although not in direct contact with it. The mutation T478K is stabilizing and predicted to decrease local protein flexibility.

### Surface and interface analysis

The entire set of mutations in position 417, 452, 478, 484 and 501 are in the Spike RBD at the interface with the ACE2 N-terminal helix (**Figure 1**). However, we note that the mutant positions 452, 478, 484 and 501 are within the RBM, containing residues that bind to ACE2. On the other hand, the mutant position 417 is located outside the motif (14).

**Figure 1.**
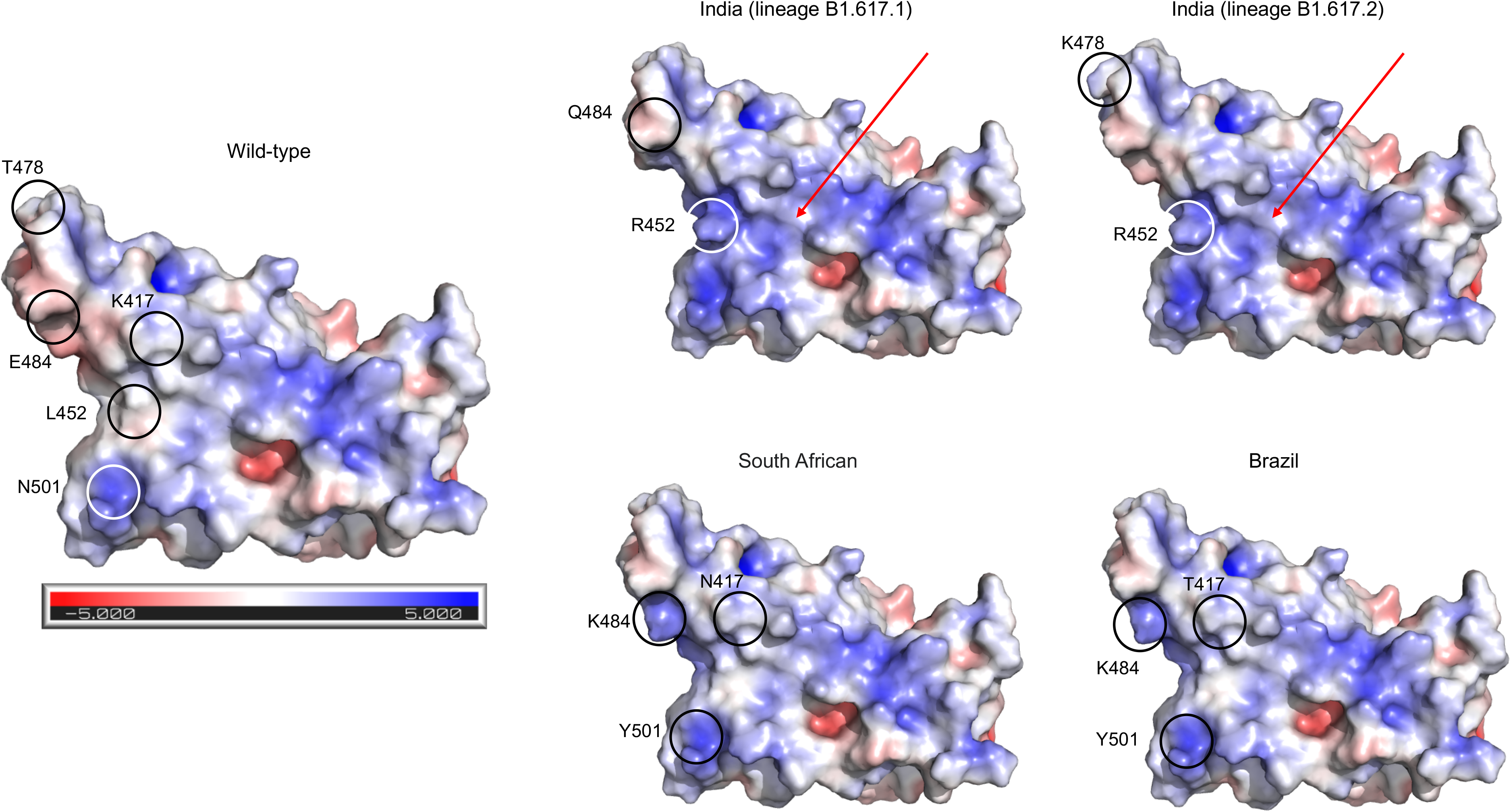
Comparison between the wild-type and variant Spike RBDs. Protein surface is coloured according to the electrostatic potential. Colour scale ranges from −5 kT/e (red) to +5kT/e (blu) as reported by the bar under the wild-type RBD. Position of the mutant sites are indicated by a circle and an attached label. Red arrows mark the area of icreased positive potential in the RBD indian variants.

According to our analysis, the mutations in position 417, 484 and 501 might alter the Spike binding affinity with the ACE2 receptor. In particular, the Tyr replacing Asn501 may form an aromatic interaction with ACE2 Tyr41, a hydrogen bond with ACE2 D38 and a potential cation-π interaction with ACE2 Lys353. In addition, substitutions of Glu484 with Lys or Gln may form hydrophobic interactions to Ile472, Gly482, Phe486, Cys488 and Tyr489. Our data also indicate that replacing Lys or Gln with Glu484 abolishes the interfacial salt bridge between Glu484 and ACE2 Lys31. Due to the fact that Lys417 is solvent-exposed and forms salt-bridge interactions with Asp30 of ACE2, replacement of Threo/Asn with Lys417 could abolish this interaction. Moreover, although the mutations in positions 452 and 478 are within the receptor binding motif, our analysis does not show a direct interaction with ACE2.

### Electrostatic potential

We note that a major, global effect of the mutations characterizing the Indian variants is represented by the alteration of the RBD surface electrostatic potential. In particular, in the lineage B.1.617.1 the uncharged and hydrophobic residue Leu452 changes to the positively charged residue Arg, and the negatively charged residue Glu484 is replaced by the uncharged residue Gln. On the other hand, the B.1.617.2 lineage shares the same mutation in position 452, but it has another mutation in position 478 where the neutral residue Thr changes to the positively charged Lys. The presence of two positively charged residues in the variant B.1.617.2 increases the positive electrostatic potential surface (**Figure 2**).

**Figure 2.**
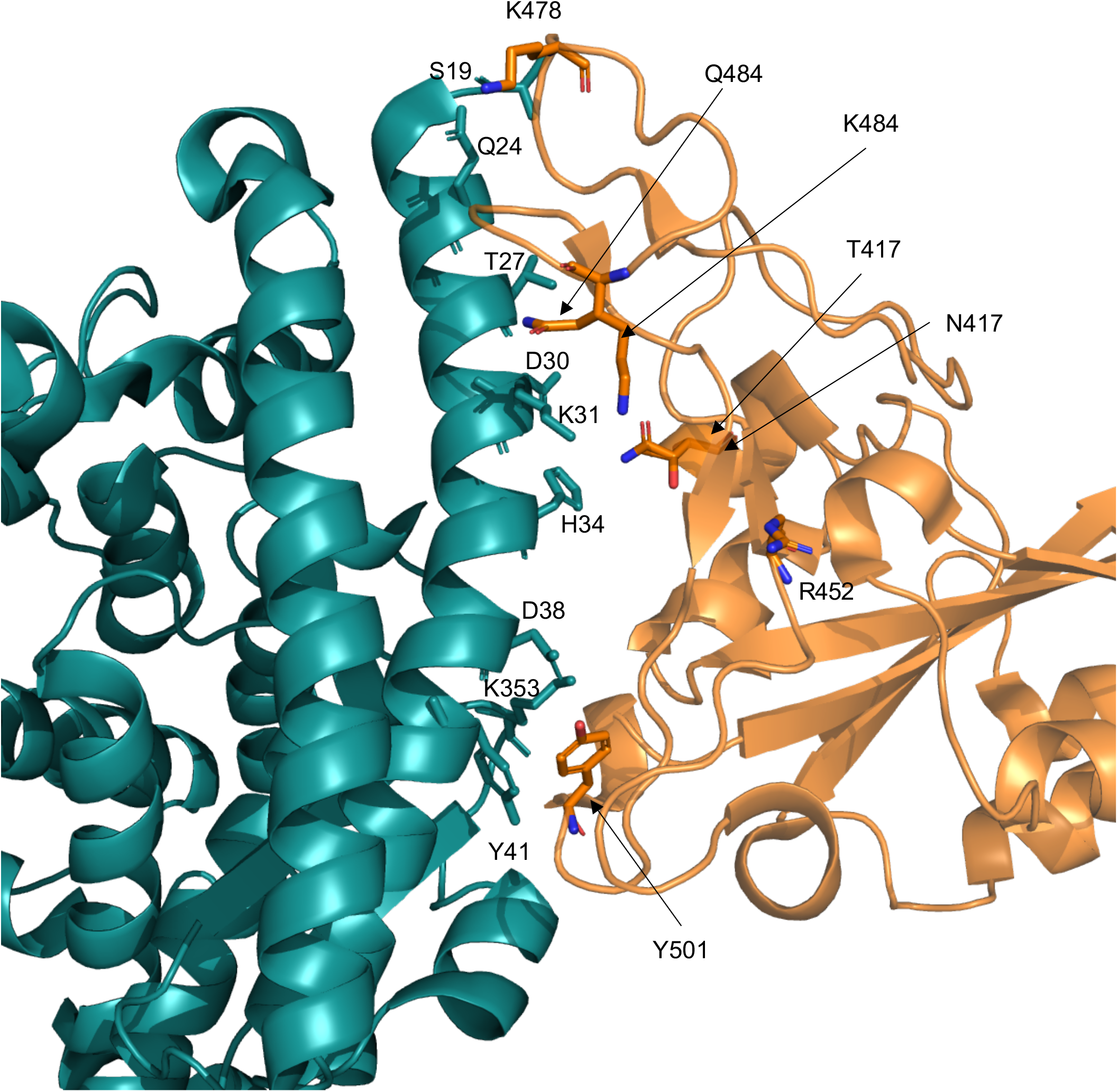
Ribbon model of the interface between ACE2 (deepteal) and RBD (orange). Side chains of relevant residues are displayed as stick models and labelled. The two mutations at the RBD sits 417 and 484 have been simultaneously displayed.

## Discussion

Given that SARS-CoV-2 variants are characterized by evolutionary and genetic changes accumulated in the genome, the use of new and improved phylodynamic techniques for the study of how epidemiological, immunological, and evolutionary processes contribute to shaping viral phylogenies is extremely important and useful. Here using a well-structured software workflow we provide evidence of a strong system able to carry out a quick, systematic and reproducible screening of the SARS-CoV-2 genome isolates. Protein mutations were identified by scanning high-quality SARS-CoV-2 genomes variants downloaded from GISAID databank (15). We reasoned that the most likely health-threatening properties of SARS-CoV-2 VOC rely on fine biochemical and biophysical changes that eventually impact the RBD interaction with the ACE2 receptor present on the host’s cell surface. We thus characterized B.1.617+ SARS-CoV-2 VUI using in-silico methods capable of predicting the effect of mutations on S-RBD protein stability and its electrostatic potential. The B.1.617+ mutations have been investigated by comparing most of the already known mutations of previously reported VOCs (16). To further justify our assumption, we note that the intensely investigated D614G substitution of the spike protein, early reported in Italian isolates (17, 18, 19), and subsequently attributed with increased virus transmissibility (20) was found to enhance the protein torsional ability and potentially affecting its stability (20).

Regarding the commonly denominated Indian variants, these constitute the new SARS-CoV-2 lineage B.1.617, which is actually composed by a family of three sub-variants, namely B.1.617.1, B.1.617.2 and B.1.617.3. This lineage, which emerged in India in October 2020 has since spread to other countries, particularly in UK, in settings with high density of residents immigrants from India. Data from Public Health England registry shows that the subvariant B.1.617.2 has become epidemiologically prevalent (21). Because of its amino-acid composition, the variant has been designed as B.1.617+ by the WHO (22).

A dichotomic behavior is observed with variant B.1.617.2. On the one hand, it lacks the mutation “E484Q” which is present in the other two lineages, and this absence seems to confer a certain degree of resistance to antibody neutralization. On the other hand, this subvariant has a mutation at the site 478 where a lysine replaces the proline. Of note, variant B.1.617.2 is indeed characterized by a major shift towards increased positive electrostatic potential because of three amino acids changes from negative or neutral to clearly positive charge, as shown in the Results section.

Sub-variants B.1.617.1 and B.1.617.3 have both the double mutations E484Q and the L452R. Although initially believed to enhance the antibody-escape potential, it has been shown that the B.1.617.1 subvariant is pretty neutralized by the majority of sera from convalescent individuals and all sera from vaccinated subjects (24). Nonetheless, this variant has been shown to be more pathogenic than the B.1.1.7 variant in an experimental model of SARS-CoV-2 infection in hamsters (25).

Our data indicate that most of the mutants are predicted to destabilize the RBD structure, except for T478K. It is thus conceivable that with destabilization comes altered binding affinity to ACE2 and to neutralizing antibodies. As noticed above, the influence of the mutations on the Spike surface properties is particularly evident in the B.1.617+ lineage, particularly for the B.1.617.2 lineage component, where the positive electrostatic surface potential is markedly increased. This may favor RBD interaction with the negatively charged ACE2 (26), which in turn would then increase affinity for the ACE2 receptor. All of these changes have the potential to eventually modify infectivity, pathogenicity and virus spread. Regarding differential binding to neutralizing antibodies, previous studies suggested that VOCs RBDs changes in the electrostatic potential surface could induce SARS-CoV-2 antibody evasion and even single amino acid changes that marginally affect ACE2 affinity could greatly influence nAbs affinity (27). Several factors have been demonstrated to affect the impact of VOCs. For example, it has been observed an increased effect at pH associated with nasal secretions (from 5.5 to 6.5) (28). For this reason, additional experiments both in vitro and in vivo are needed to establish the biological significance of SARS-CoV-2 mutations and how the interactions between mutations and local cellular microenvironment influences the clinical outcome and the transmission dynamics of this virus.

## Acknowledgments

SP has been partially supported by the Sapienza grant RP120172B49BE24. Authors are grateful to Alice Vinciguerra for her precious assistance. We also would like to thank all the authors who have kindly deposited and shared genome data on GISAID.

